# Spectral memory: Illumination-evoked plasticity

**DOI:** 10.1101/490888

**Authors:** Ifedayo-Emmanuel Adeyefa-Olasupo

## Abstract

Here human trichromats were presented with two types of scenes – geometric and real-world scenes –tinted with a shade of colour in order to destabilize the perceived illumination, and chromaticity of the retinal image of each scene. Each trichromat was instructed to adjust the chromaticity of the object embedded within each scene until its surface appeared devoid of any hue in the DKL colour space which spans two chromatic opponent axes – the S–(L+M) and L– M axis – and a luminance axis – the L+M axis. The following observations were made : (i) across scenes, adjustments were dispersed along the S–(L+M) axis, along which daylight is known to vary; (ii) across trichromats, for the geometric scenes, adjustments were biased towards the S pole of the S–(L+M) axis for one group (group 1), and towards the (L+M) pole for the other group (group 2); (iii) for the real-world scenes, adjustments for both groups systematically converged towards the (L+M) pole. These results suggest that when the core set of priors upon which the human visual system typically relies become ill-equipped, the human visual system is able to recruit one of the two illumination priors – PriorS or PriorL+M – in combination with the representation it has formed over time about the spectral composition of the illuminant associated with scenes the trichromatic observer is currently being exposed to within its ecological niche, as it attempts to stabilize the chromaticity of the retinal image of real-world scenes.

## Introduction

An array of distal properties contributes to the proximal retinal image of a scene – the primary, secondary, and tertiary illuminant, the transmittance properties, the reflectance properties of objects within the scene, the organism’s relative viewing orientation and the optics of its eye, along with other factors which actively modulate the incoming retinal signal. This signal, however, is insufficient with respect to the inverse problem of perception, that is inferring the components of these distal properties from the retinal image. To this end, the perception relies on, not one, but a combination of implicit assumptions and inferences (or internal priors) [1,2], many of which have been strongly influenced by the organism’s past and current ecological niche [3,4,5].

Consider Figure 1A. Here a simplified snapshot of a poorly illuminated retinal image of a scene at an early stage in visual processing is given. If visual processing were to terminate here, the normal trichromatic observer (vision which is based on L, M, and S cone photoreceptors) will most likely perceive an unstable retinal image, that is, a scene, composed of geometrical objects, which appears to have three sets of colours: yellow and white, blue and brown, and blue and black. This ambiguity however does not make for an efficient visual system, especially in cases when the chromaticity of the retinal image is critical in assessing the state of an object (e.g. ripe or unripe fruit, green or red traffic light) embedded within a scene or enhances the recognition of an ambiguious scene [6]. Aware of this erroneous representation of the retinal image, the visual system must undergo the appropriate type of operation in which the spectral components, namely the chromaticity which makes up the illuminant source, is calibrated, thus stabilizing the retinal image of the scene [7]. However, in conditions like Figure 1A, the canonical priors and operation the visual system usually undergoes (e.g. chromatic adaptation), in which the chromaticity of the incident illumination discounted, for the most part, is consistent across normal trichromats, does not always hold, as it is observed in the dress phenomenon [8,9]. Specifically, when normal trichromats are instructed to report the colours perceived in Figure 1A, a great deal of individual differences is observed, that is, the retinal image is systematically reported as appearing in either yellow and white – 1B, blue and black-1C, or brown and blue-1D, responses which suggest that the chromaticity being discounted by the human visual system across trichromats is radically inconsistent [10,11].

**Figure 1.**
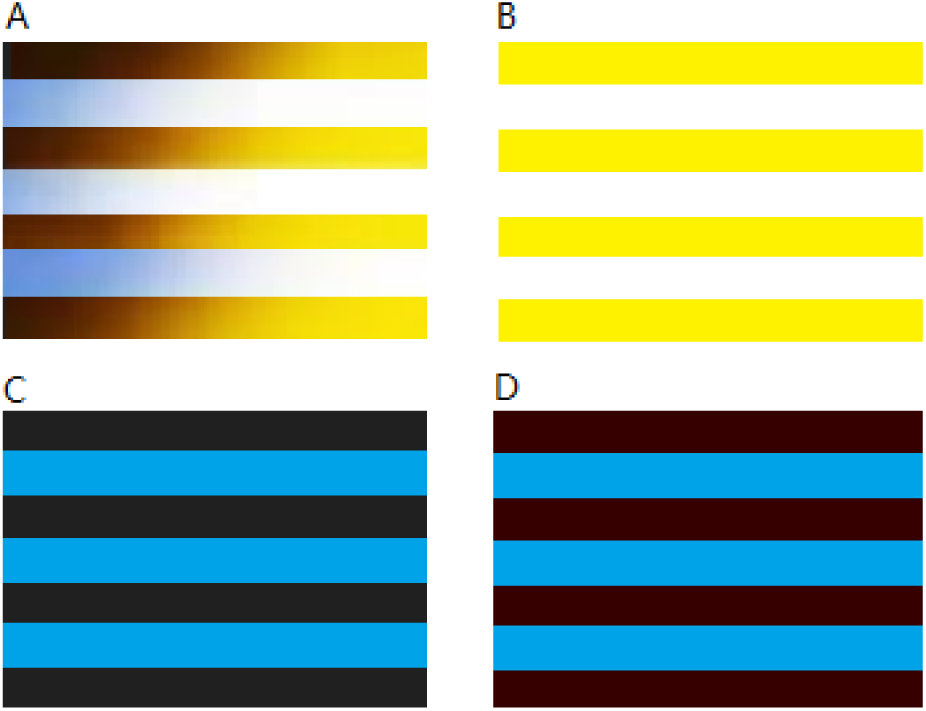
Striking individual differences in perception. A. an uncalibrated retinal image under poorly illuminated conditions, percept the visual system constructs when it elects to discount the short-wave illuminant – B, long-wave illuminant – C, or both illuminant – D.

**Figure 2.**
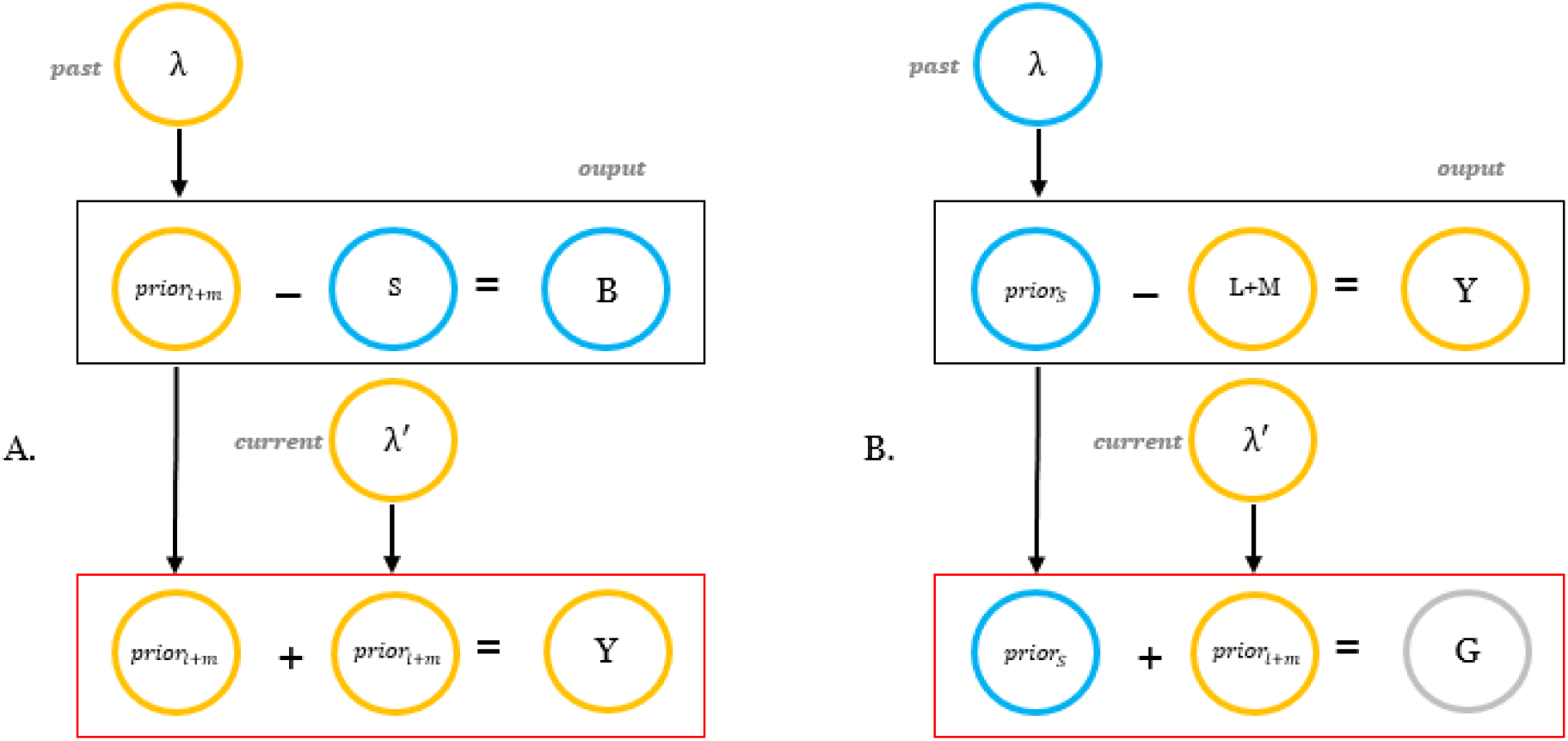
The effects of spectral memory on chromatic perception. A. group 1 dynamics, B. group 2 dynamics. Depending on the illumination the visual system’s most basic architecture was preferentially tuned under, during its development – λ, an illumination prior emerges and is stored – PriorL+M or PriorS. Because the mathematical operation employed by the visual system for the geometric scenes-the first rectangular box, is subtractive in nature, perception will most likely shift towards that prior’s opponent color. For the real-world scenes, λ′ denotes the chromaticity of the illuminant scenes observed within the trichromat’s current ecological niche is being illuminated by. Since the operation employed by the visual system for these class of scenes-the red rectangular box, is additive, both priors – the developmental and empirical prior combined (i.e. spectral memory) go on to modulate the perception of real-world scenes [see Figure 5A and 5B].

To explicate this divergence in perception, recent studies have put forth an incredibly novel and compelling hypothesis [12,13]. Considering that the mammalian visual system, specifically the molecular machinery within the retina (i.e. rhodopsin molecules) which enables the detection of photons of light, as well as the thalamic and cortical circuits which subserve visual perception (e.g. lateral geniculate nuclei and striate cortex) developed under daylight illumination which routinely fluctuates from shorter wavelength of light in the morning towards longer wavelength of light during the course of the day, the mammalian visual system (including humans) may possess one of two internal priors – *PriorS* (blue) or *PriorL+M* (yellow) which project along the S–(L+M) (blue-yellow) axis, along which daylight illumination is known to vary [14]. Consequently, these internal priors, derived from the organism’s past ecological niche, under ill-defined conditions, according to these authors, can be used to stabilize the retinal image of a scene. This means that, in cases like Figure 1A, the visual system which is preferentially tuned to short-wave illuminants (e.g. blue skies) expects the spectral composition of the illuminant to be largely composed of the bluish spectrum, which leads it to discount the bluish component of the illuminant, thus constructing a whitish and yellowish percept, that is, a projection towards the L+M pole (yellow) of the S–(L+M) axis. Inversely, for the system preferentially tuned to long-wave illuminants (e.g. direct sunlight) the yellowish spectrum of the illuminant is quickly discounted, resulting in a blackish and bluish percept, that is, a projection towards the S pole (blue) of the S–(L+M) axis. Finally, in the most extreme case, the visual system tuned to both illuminant types, discounts both the bluish and yellowish spectrum of the illuminant, resulting in a brownish and bluish percept.

While this hypothesis appears plausible, the jury is still out as to whether these internal illumination priors are in fact represented on the perceptual domain of the human visual system. Specifically, it is not known whether the visual system is only capable of possessing one of the two internal priors – PriorS or PriorL+M, both, or combines different types of priors, as it is observed under less ill-defined conditions. One way to empirically address this question is to measure the chromatic perception of real-world scenes, many of which the human trichromat is currently being exposed to versus scenes (geometrical scenes) which are not observed within its current ecological niche, with the illumination under which they are presented being equally ill-defined – so as to examine whether different types of priors are recruited for different types of scenes. Assuming that the scene-type does not matters, then the visual system whose basic architecture is preferentially tuned to long-wave illuminants (direct sunlight), for both real-world and geometrical scenes, will naturally discount the yellowish spectrum of the illuminant, thereby shifting the chromaticity of the retinal image towards a bluish percept. Inversely for the visual system whose architecture is preferentially tuned to short-wave illuminants (bluish skies), a yellowish bias should be observed for both scenes. Alternatively, if scene-type does in fact matters, then it is possible that the visual system which has developed under long-wave illuminated scenes, when presented with a real-world scene, should naturally discount the yellowish spectrum of the illuminant, thereby shifting the perception of the scene towards a bluish percept. Inversely, for the opposing class of scenes– the geometric scenes–, considering that these scenes do not approximate conditions under which that particular visual system developed under, the illumination prior recruited in stabilizing the chromaticity of these scenes, in the most extreme case, may be opposed to the observed primary illumination prior. Under this same assumption, the opposite results would also be expected for the system which developed under short-wave illuminated scenes. Finally, another possibility could be that a more empirically derived illumination prior (the mean chromaticity of the illuminant), one which is based on the trichromat’s current ecological niche, is used in combination with PriorS or PriorL+M, to help provide a stable retinal image.

To foreshadow the results in the experiments which are reported here, when normal trichromats were instructed to adjust the surface of a set of natural fruit objects or a circular disc, which are embedded within a scene, until it appeared devoid of any hue, chromatic adjustments were almost always projected along the S–(L+M) opponent axis, along which daylight is known to vary. Specifically, two groups emerged. The first group of trichromats (group 1) systematically adjusted the surface of the circular disc (the geometrical scenes) to the (S) bluish pole of the S–(L+M) opponent axis, while the natural fruit objects (the real-world scenes) were adjusted towards the L+M pole of the S–(L+M) opponent axis – a remarkable demonstration of rapid visual plasticity in the adult visual system [15]. As for the second group (group 2), chromatic adjustments for the geometrical scenes were projected towards the (L+M) of S–(L+M) opponent axis. And while the degree of bias towards the S pole was significant for individual trichromats, overall chromatic adjustments across trichromats were almost perfectly aligned with the origin of the DKL color space for the real-world scenes. These results suggest that the human visual system possesses a core internal illumination, which it principally recruits under ill-defined conditions, as it is reflected in the chromatic adjustments for the geometrical scenes. Results further suggest that under these same illuminant conditions, for the real-world scenes, for both groups, on average, this developmentally derived prior – PriorS or PriorL+M, is combined (or in Bayesian terms updated) with a more empirically derived prior, to form what I have termed “spectral memory” – an additive chromatic representation of the illuminant.

Consistent with previous reports under normal conditions [16,17], it is the combination of these two priors the human visual system appears to recruit, as it attempts to stabilize the chromaticity of the retinal image of real-world scenes under extremely ill-defined conditions [1,2, see Figure.2]

## Method

### Human trichromats

Twenty-five naive normal human trichromats concurred to participate in part 1 and part 2 of the experiment. There was a total of fifteen females and ten males with a mean age of 23.11 and an SD of 3.98. Each trichromat was tested for normal to corrected-to-normal visual acuity and normal colour vision using the Ishihara colour plates. The protocol for this study was approved by the Johns Hopkins University Homewood IRB.

### Apparatus

Stimuli were presented on a Dell D1526HT cathode ray tube (CRT). Gamma correction and colour space calibration (computed using spectral distribution of the CRT’s phosphor channel and the Smith & Pokorny 2-deg cone fundamentals) were accomplished using standard procedures [18,19] and utilizing a UDT Instruments model 370 optometer with a model 265 photometric filter (mounted atop a Dolica 68” Lightweight Tripod) and a Lighting Passport Pro spectroradiometer. The CRT was firmly positioned 30cm behind a neutral wall which subtended 81 × 42°, allowing the concealment of the light source. CRT was driven by a NVIDIA Quadro FX 580 graphics card with a colour resolution of 8 bits per channel. The spatial resolution of the CRT during the duration of the experiment was 1280 × 1024 pixels with a refresh rate of 85 Hz (11.76 ms per frame). The Judd correction CIE xyY values for the red, green and blue primary were (x=0.631 y=0.337 Y=20.451) (x=0.295 y=0.600 Y=52.743) (x=0.148 y=0.065 Y=6.457) respectively. Directly at the center of the wall was an opening which subtended 10.6 × 9° [20]. The interior region of the viewing tunnel was homogenously painted using a Clarke and Kensington deep matte black acrylic paint, thereby inhibiting any unsolicited shadows or light reflectance. This was later connected to the posterior region of the neutral wall directly to the center of the CRT’s phosphorescent surface. At the anterior base of the wall extended 17cm horizontally a trapezoid-shaped hand chamber with the exterior and interior painted deep matte black to prevent any reflectance of keypress or any glossy or self-illuminating objects worn by trichromats. Inside the hand chamber was fixedly placed a black Dell L100 keypress. Attached to the base of the hand chamber was a black adjustable head and chin rest. The chin rest not only allowed for a standardized viewing distance across trichromats at 52 cm, but also allowed for a natural demarcation between the left and right-hand. Figure 3A-B presents a photograph of the experimental setup.

**Figure 3.**
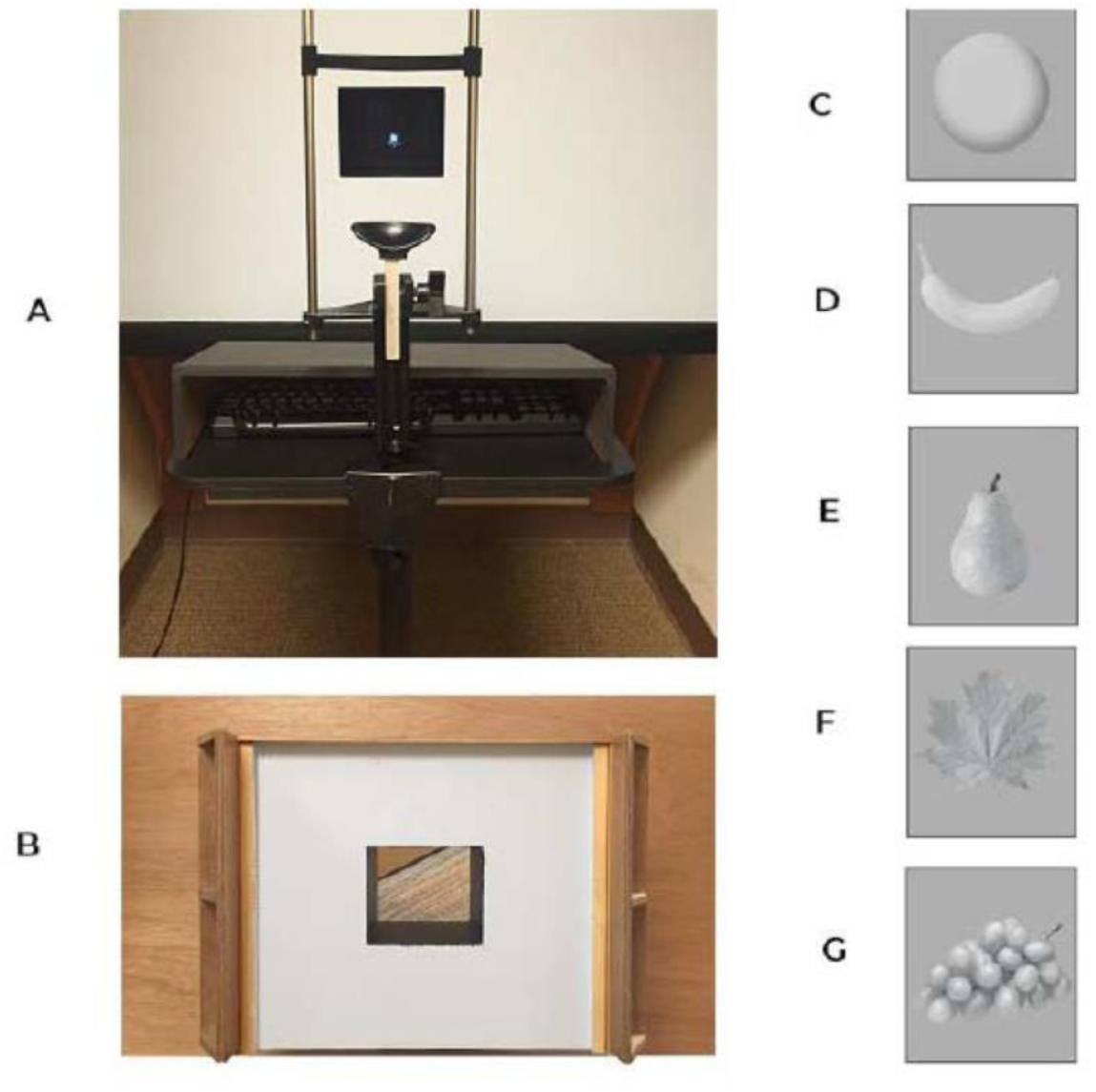
Photograph of the apparatus and stimuli (resized for viewing purposes) used in the present study. (A) trichromat’s view during the experiment; (B) the dorsal view of the viewing tunnel which was firmly placed on the front of the CRT; (C) example of a circular disc; (D) banana; (E) pear; (F) foliage; (G) grapes.

### Stimuli & Procedure

In each trial of part 1 of the experiment, each trichromat observed a homogenously coloured circular disc (3×3°). This object was carefully selected as it is an object which approximates real-world objects to a lesser degree. The luminance of the disc was set to 30 cd/m^2^ while the background was set to 28.632 cd/m^2^ in order to avoid any colour blending strategies [21]. The circular disc appeared with a tinted shade of colour chosen randomly from within the DKL colour space at a radius of 0.10 (10% of the maximum saturation allowed in the DKL colour space). The shade was created by digitally manipulating the pixels of each object (see Figure 3) in order to create a shadow-like casting on the surface of the object, that is, make it ambiguous, such that at the onset of each trial the trichromat became less certain about the illuminant source and consequently the achromaticity of the object [22,23]. Note that the colour space used to define the stimuli in the experiment reported here is modeled after the second stage of colour perception, thereby reflecting the preferences of neurons in the Lateral Geniculate Nuclei (LGN) non-human trichromats (rhesus macaques) [24,25]. Furthermore, the trichromats’ task was to adjust the hue of the disc until it appeared void of any hue. Adjustments were made by the keypress, along the isoluminant L–M (reddish-greenish) colour opponent axis, using the left arrow and right arrow, or along the isoluminant S–(L+M) (bluish-yellowish) axis, using the up and down arrows. Trichromats were also able to modify the pace by which they preferred to move along each axis, toggling between finer and coarser changes by pressing the ‘v’ and the ‘t’ keys respectively. If a given trichromat got lost within the colour space, he or she could press the ‘q’ key, resetting the object to the initial colour it was presented in. Finally, upon the completion of an adjustment (pressing the ‘a’ key), the chromaticity of each individual pixel was scaled to the stimulus’ new chromatic position [19,21]. While there were no time constraints on the adjustments, the adjustment time was recorded. In the event a given trichromat pressed the “q” button the adjustment time was set back to its initial starting point.

Trials in part 2 of the experiment were identical to trials in part 1, with two critical exceptions. Firstly, each trial included one of four digitalized photographs of natural fruit objects captured under real-world conditions, instead of a circular disc. The banana was subtended 6.20° × 4.07°, the grapes, and the foliage was subtended 4.66° × 4.57° and the pear was subtended 4.41° × 5.56°. These objects were set to have the same mean luminance values across their surfaces as the background (28.632 cd/m^2^). Secondly, in other to intensify the uncertainty of the illuminant source, upon the first appearance in a trial, these objects did not appear in a random hue. Instead, the surface of each object was covertly tinted a canonical, deuterocanonical or opponent hue associated with the object. This was done by digitally manipulating the pixels of each object at a saturation level of 10%. In the ‘canonical’ condition, an object appeared with a tint of a canonical hue (e.g. purplish tint in the case of grapes, see Figure 4B). In the ‘deuterocanonical’ condition, an object appeared in a tint of a hue of a secondary association (e.g. yellowish for a pear). In the ‘opponent’ condition, the object was tinted with a hue situated in the opponent direction from its canonical association (e.g. blueish in the case of a banana). Although I call them ‘part 1’ and ‘part 2’ their order was counterbalanced in practice across trichromats. To maximize the reliability of mean chromatic adjustment value, each trichromat completed a total of 8 trials in part 1 and 80 trials in part 2 (equally distributed across objects and conditions).

**Figure 4.**
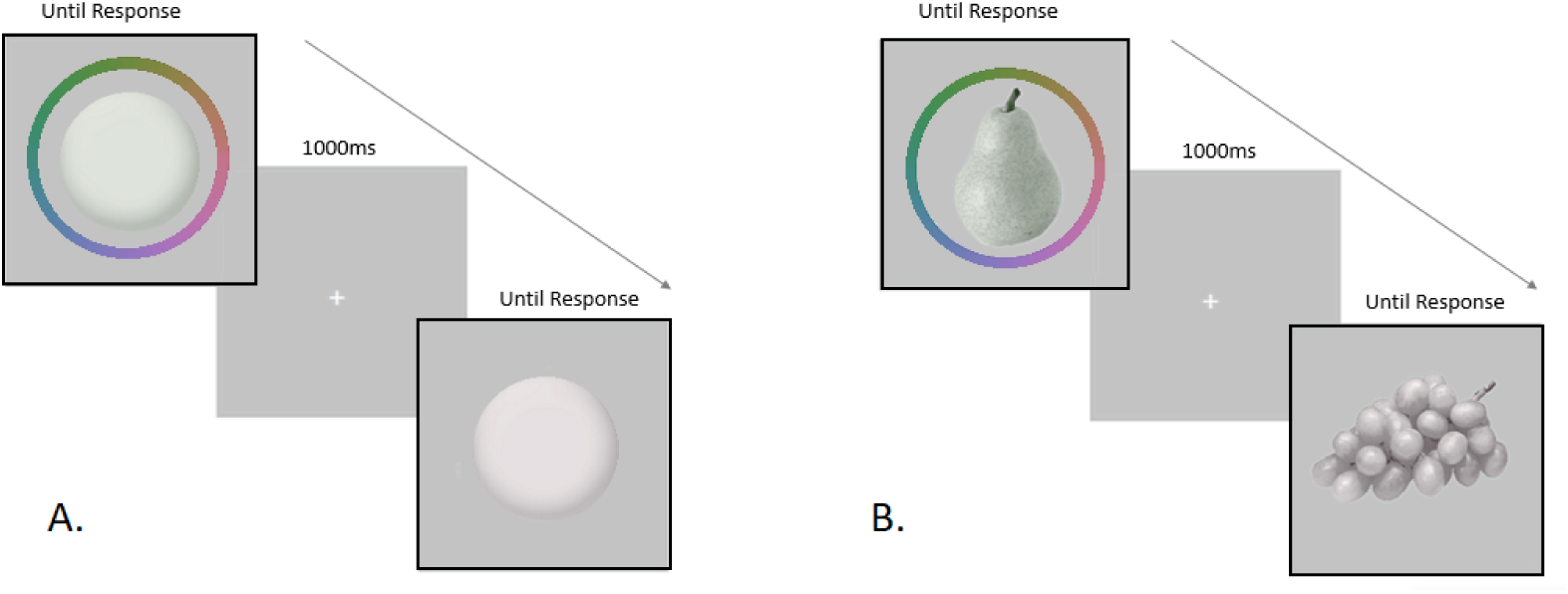
General procedure for part 1 and part 2: (A) circular disc, (B) natural fruit objects tinted in a greenish or purplish hue in order to create a shadow-like casting on the surface of the object. The colour ring (for illustration purposes) around the first trial denotes the possible colours within the DKL color space trichromats could adjust the surface of the object to. The 1000ms fixation window which follows each trial was intended to control for any afterimage effect following the offset of each trial. The next trial without the color ring is a more accurate representation of what each human trichromat actually saw at the onset of each trial.

## Results

### Part 1 – geometrical scenes

Using the horizontal meridian as the chromatic boundary to examine whether two types of groups would emerge, one shifted towards the S pole of the S–(L+M) axis, and the other shifted towards the (L+M) pole of this axis, each trichromat’s mean chromatic adjustment was computed in part 1 across all trials. Shown in Figure 5A, the chromatic mean (blue diamond) revealed that 14 (females=9, males=5) out of the 25 trichromats adjusted the circular disc towards the S (bluish) pole to varying degrees. The mean age for this group was 20.28 with an SD of 2.78. The mean chromatic adjustment along the x-axis across all trichromats was −0.0999 with SEM of 0.0344, while along the y-axis the mean chromaticity was −0.0785 with a SEM of 0.0257. For the remaining 11 (females =6, males =5) trichromats, the mean chromatic adjustments (yellow diamond) shown in Figure 5B, were biased towards the L+M (yellowish) pole of the S–(L+M) axis, while the mean age was 26.27 with SD of 2.24. Here, the mean chromaticity along the x-axis was 0.0062 with SEM of 0.0138, while along the y-axis, the mean chromaticity was 0.0599 with SEM of 0.0135.

**Figure 5.**
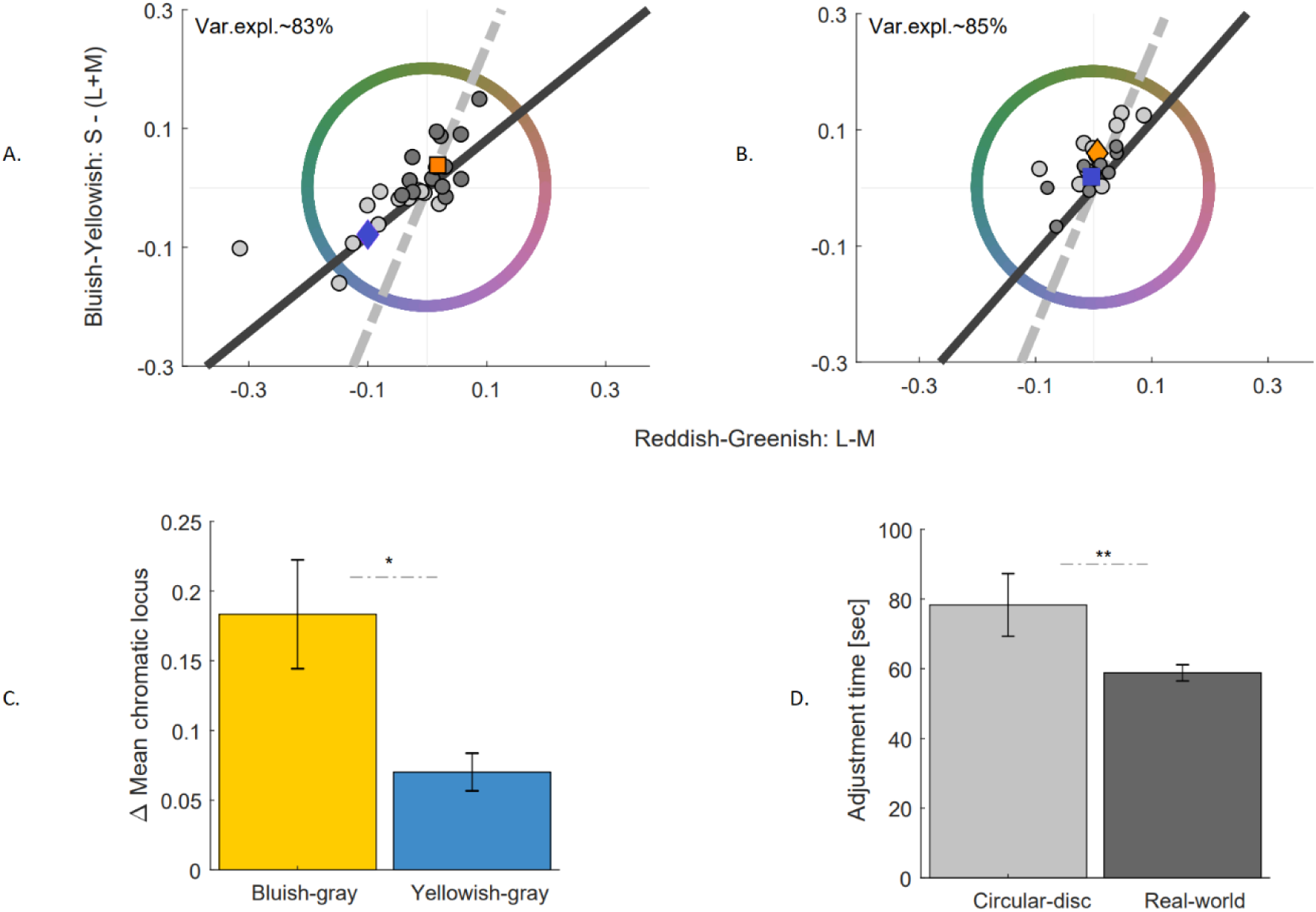
(A)Mean chromatic adjustments for the circular disc and natural fruit objects for group 1(n=14). (B)Mean chromatic adjustments for the circular disc and natural fruit objects for group 2(n=11). Light gray circle denotes the mean chromatic adjustments of the disc for each trichromat within each group, while the dark gray circle denotes the mean chromatic adjustments of the fruit objects. Mean chromatic adjustments across trichromats in each group for the disc and fruit objects is denoted with a diamond and square symbol respectively, while the filled color for each symbol indicates the color in DKL color space the mean chromaticity is trending towards. The black solid line is the 1^st^ principal component axis, and the dotted gray line denotes the daylight axis. (C) Change, in Euclidian units, the mean chromaticity between the disc and the fruit objects for group 1 (yellow bar) and group 2 (blue bar) with their associated standard error of the mean. (D) Adjustment time across groups for the disc (light gray bar) and natural fruits (dark gray bar) with their associated SEM.

### Part 2 – real-world scenes

In the natural fruit conditions – part 2, similar to my observation in part 1, the mean chromatic adjustments showed two distinct patterns, those whose chromatic adjustments were biased towards the yellowish pole of the S–(L+M) opponent axis and those whose chromatic adjustments were trending towards the bluish pole of the S–(L+M) opponent axis. What is of particular interest is that trichromats, to varying degrees, systematically adjusted the chromaticity for these objects towards the opponent direction of their mean chromatic locus in part 1. Specifically, for those whose mean chromatic adjustments were biased towards the bluish pole of the S–(L + M) opponent axis in the case of the disc, for natural fruit objects – irrespective of the canonical, deuterocanonical or opponent hue in which they were tinted – average chromatic adjustments were projected towards the yellowish pole of the S–(L+M) opponent axis (see Figure 5A, yellow square). The mean chromatic adjustments across all objects along the x-axis was with SEM of 0.0099, while the mean chromatic adjustments along the y-axis was 0.0384 with SEM of 0.0131. For the group of trichromats whose mean chromatic adjustments was projected towards the L+M pole in part 1, adjustments for the natural fruit objects trended towards the opponent direction, that is, towards the S pole when compared to the adjustments for part 1 (see Figure 5B, blue square). Across all trichromats, the mean chromatic adjustments along the x-axis was −0.0044 with SEM of 0.0114, while the mean along the y-axis was 0.0185 with SEM of 0.0111.

### Variation along the S–(L+M) axis

For each group, in order to assess whether part 1 and part 2 chromatic adjustments covaries along the S– (L+M) opponent axis, and to what extent adjustments were distributed along the daylight axis (gray dotted line), a principal component analysis (PCA) was performed [26]. For the first group, the primary principal component axis explained 83% of the observed variance, while the direction of the main covariation was primarily projected along the S–(L+M) opponent axis. For the second group, 85% of the variance is explained by the primary principal component axis, which was also projected along the S–(L+M) opponent axis. For both groups, the principal component axis aligned much better with the daylight axis along the L+M pole of the S–(L+M) opponent axis in comparison to the chromatic adjustments around the S pole.

### Difference in distributions between scenes

To investigate whether the chromatic distribution within group is statistically significant between the disc and natural fruit objects, a two-sample Kolmogorov-Smirnov test was performed [27]. For the difference between the first group, distributions along the x and y-axis for part 2 was significantly different from normality, that is, the distribution observed in part 1 (x-axis and y-axis, p<0.01). For the second group however, the distribution in part 2 was not significantly different from normality (x-axis, p= 0.7358 and y-axis, p=0.1473).

### Spatial and temporal dynamics

Chromatic displacement for each trichromat was computed by taking the Euclidean distance between the mean chromatic adjustment in part 2 from the mean chromatic adjustment in part 1. Following this, the mean chromatic displacement across trichromats within each group was computed (see Figure 5C). For the group whose mean was projected towards the S pole, an average displacement of 0.183 (SEM=0.03) towards the L+M pole was observed. As it is expected for the second group, a much more modest chromatic displacement of 0.070 (SEM=0.01) towards the S pole was observed. A t-test was conducted which confirmed that the observed difference between the two groups was significant (p <0.05). To assess the temporal dynamics which accompanied the observed chromatic displacement, the mean adjustment time across each group for the circular disc and natural fruit objects was computed. Surprisingly a temporal difference was observed. Specifically, results show that for the circular disc, trichromats on average took 78.30 secs (SEM=8.99) to adjust the surface of these class of objects. Inversely, a much lesser time – 58.82 secs (SEM=2.32) was required for the natural fruit objects. A t-test was performed, which confirmed the significance of this difference with p<0.01.

## Discussion

The hallmark of any dynamic system lies in its ability to rapidly modify its most basic architecture in response to external demands. Here, I sought to extend the research on how the human visual system stabilizes a visual scene in which the incident illuminant is ill-defined and study its effects on the chromaticity of the objects embedded within that scene, particularly in the cases when the core set of implicit assumptions and inferences the visual system typically relies upon become ill-equipped. While more recent studies have suggested that the human visual system deals with these cases by recruiting strictly one of two illumination priors – *PriorS* or *PriorL+M* – I investigated whether the human visual system is hard-wired to a single illumination prior – *PriorS* or *PriorL+M* – or is in fact plastic, that is, able to recruit either both types of priors or combine different types of priors to form a retinal image.

In part 1 of the experiment, trichromats systemically adjusted the surface of the circular embedded within a scene, either towards the S or L+M pole along the S–(L+M) axis, which daylight is known to vary along. These results support the general hypothesis that when the human visual system encounters a geometric scene in which the illumination is ill-defined, internal illumination priors direct the visual system as to the chromatic property of the illuminant source it is to quickly discount [10]. Secondly, these results are also consistent with the idea that trichromats, for these scenes, typically fall into two distinct groups – blue and yellow group, suggesting that different visual systems use different priors to help stabilize the chromaticity of the retinal image of a geometric scene [9,28].

In part 2, for the trichromats, who exhibited a particular bias towards the S pole, for the geometric scene, a significant opponent bias for the real-world scenes was observed, while trichromats, who exhibited a particular bias towards the L+M pole for the geometric scenes demonstrated modest opponent effects for the real-world scenes. Prima facie these results appear to suggest that the human visual system is in fact able to switch between both types of illumination prior – *PriorS* or *PriorL+M*. However, this explanation does not hold if the modest effects observed for group 2 is considered. Alternatively, if we consider the fact that (i.) most of the scenes human trichromats observe today, particularly those which include natural fruits objects, are principally illuminated under artificial yellowish illuminants (incandescent bulbs) [16], and (ii.) that perception relies on not one, but a combination of priors [1,2], then it appears that chromatic adjustments observed in part 2, are actively being modulated by a combination of either *PriorS* or *PriorL+M*, and the mean chromaticity of illuminant illuminating the scenes which current populate the human trichromat’s current ecological niche. This summed prior, as a result, biased group 1’s adjustments towards the L+M pole of the S–(L+M) axis, and group 2’s adjustments in and around the center of the S–(L+M) axis. This is a particularly plausible explanation if another line of evidence is considered – the difference in adjustment times for each scene. Keeping mind that more computational energy is required for subtractive operations compared to additive operations [29], it is particularly telling that for the geometric scenes, which I argue the human visual system undergoes a subtractive operation, trichromats took much longer to decide whether the surface of the disc was devoid of any hue. Inversely adjustment times on average for the natural fruit objects, which I argue is operationally additive, occured within a much shorter temporal window [Figure 5D].

In all, previous studies on scene statistics have vigorously debated the type of spectral information the visual system stores when a scene is being observed, and how this information is later recruited by the visual system to help stabilize the chromaticity of the retinal image of a scene during future viewing conditions. [30,2]. Here I argue and demonstrate for the first time that human visual system possesses a core illumination prior, which is developmentally derived. This prior is then updated (or combined) with illumination priors which are derived from the human trichromat’s day to day illumination experiences within its current ecological niche. It is ultimately the sum of these two types of illumination priors (spectral memory) which directly modulate how the human visual system stabilizes the chromaticity of the retinal image of a scene. To further assess the validity of this idea, future studies should focus on real-world scene which are principally illuminated under different types of illuminant, and how in combination with the visual system’s core set of priors – *PriorS* or *PriorL+M*, help shape the chromaticity of the retinal image of a scene. For example, one way to test this idea is to measure real-world scenes which are typically illuminated under bluish illuminants (bluish skies). To this end, one would expect that those in group 1 who are measured in this study, assuming they were asked to adjust the chromaticity of objects (e.g. sand) embedded within these types of scenes, would demonstrate a gray bias, while those in group 2 would demonstrate a blue bias. In conclusions, the scene composition and the chromatic adjustment procedure developed for this study now appears to be a useful tool for further research on illumination priors, and more generally, for investigating possible interactions between priors and its modulatory effects on the retinal image of a scene.

